# BioModelos: a collaborative online system to map species distributions

**DOI:** 10.1101/432617

**Authors:** Jorge Velásquez-Tibatá, María H. Olaya-Rodríguez, Daniel López-Lozano, César Gutiérrez, Iván González, y María C. Londoño-Murcia

**Author notes:** Current address: Facultad de Ciencias Biológicas y Aplicadas, Universidad Militar Nueva Granada, Campus Nueva Granada, Cajicá, Colombia (Jorge Velásquez-Tibatá).

## Abstract

Information on species distribution is recognized as a crucial input for biodiversity conservation and management. To that end, considerable resources have been dedicated towards increasing the quantity and availability of species occurrence data, boosting their use in species distribution modeling and online platforms for their dissemination. Currently, those platforms face the challenge of bringing biology into modeling by making informed decisions that result in meaningful models. Here we describe BioModelos, a modeling approach supported by an online system and a core team, whereby a network of experts contributes to the development of species distribution models by assessing the quality of occurrence data, identifying potentially limiting environmental variables, establishing species’ accessible areas and validating qualitatively modeling predictions. Models developed through BioModelos become publicly available once validated by experts, furthering their use in conservation applications. This approach has been implemented in Colombia since 2013 and it currently consist of a network of nearly 500 experts that collaboratively contribute to enhance the knowledge on the distribution of a growing number of species and where it has aided the development of several decision support products such as national risk assessments and biodiversity compensation manuals. BioModelos is an example of operationalization of an essential biodiversity variable at a national level through the implementation of a research infrastructure that enhances the value of open access species data.

## Introduction

Species distributions are an essential biodiversity variable [1,2], critical to assess species’ conservation status and trends [3,4], assess biodiversity change [5–7], guide conservation and management at the species and community levels [8] as well as to assess their ecosystem services [9], potential impacts on human activities [10,11] and health [12,13]. They also are key inputs for the calculation of indicators to evaluate countries’ progress towards achieving international targets, such as CBD’s Aichi targets [14] or the United Nations’ Sustainable Development Goals [15]. Thus platforms that consolidate and facilitate access to the highest quality data on species distributions are necessary to coordinate biodiversity observation delivery as EBV’s and aid biodiversity conservation and management globally [16].

Considerable international efforts and resources have been designated at the mobilization of primary biodiversity data (PBD), particularly through the global biodiversity information facility (GBIF). These data are fundamental for many conservation analyses based on species distributions. However, for most areas in the world, our knowledge on species distributions based on PBD is geographically biased and incomplete [17]. This situation is particularly dire in biodiversity hotspots which lack sufficient information on species distributions based on PBD even at coarse spatial scales [17–19]. Therefore, for most regions on earth, methods that generalize occurrences to areas representing species distributions are necessary to use PBD in conservation applications.

Species distribution modeling has emerged in the last two decades as a set of methods and practices to estimate species distributions [20–22]. They are based on PBD and environmental data and use a variety of statistical methods to infer the probability of occurrence or suitability in unsampled sites. As such, they are a powerful tool to overcome the wallacean shortfall [i.e. the lack of knowledge on species geographic distributions; 23], due to their ability to produce reasonable predictions with few occurrences [24,25], repeatability and ease of update. However, their implementation is not straightforward [26,27] and fully automated, large scale modeling procedures face several challenges [28,29], namely the need of expert’s knowledge to detect and correct certain types of errors in PBD [30]; select meaningful environmental covariates [31]; determine each species’ accessible area [32,33] and judge the biological realism of predictions [34].

Current platforms that provide maps of species’ distributions are either based largely on expert maps (e.g. map of life, www.mol.org) or use fully automated modeling workflows (e.g. BIEN, http://biendata.org/). Expert maps in some cases may be the only way to characterize a species’ distribution, for example when there are very few observations available (< 5), but are difficult to update as new observations accumulate [35] are not repeatable and their precision is often too coarse to inform conservation at regional scales [36]. On the other hand, large scale, fully automated modeling workflows are unable to detect and fix errors that require domain specific expertise [e.g. species misidentification or geographic outlier detection; 30] and unspecific modeling choices, for example of accessible areas and environmental variables, is likely to result in biologically unrealistic models.

To address the challenge of mapping large numbers of species without compromising biological realism we devised BioModelos (biomodelos.humboldt.org.co), an online system that involves a network of experts and a core team of modelers in the development and validation of species distribution models which are freely available for public visualization and download. Here we describe the operational approach of BioModelos, the functionalities of its web app and its implementation in Colombia. Although BioModelos has thus far been deployed in a single country, our philosophy and software architecture can may be applied to other regions and even scaled up to global implementations.

## Network structure and governance

The aim of BioModelos is to provide distribution maps for a set of species in a particular area that are validated in terms of their biological realism by experts. To that end, in BioModelos experts are arranged into groups according to their areas of taxonomic/geographic expertise. Experts are defined as individuals whom are able to either curate and improve the taxonomic and/or geographic quality of occurrence data, inform the selection of certain modeling parameters (e.g. accessible area) or assess the performance of competing species distribution hypothesis, for at least one species in their group.

Each group is coordinated by one or more moderators, whom are ideally a well-connected members of a community of researchers interested in advancing the knowledge on the distribution of a set of species. As such, they are responsible for the objective evaluation of the expertise of potential members, setting deadlines for each step in the model development workflow in agreement with group experts and expedite the completion of the group modeling agenda.

The core team of BioModelos, facilitates some of the steps in the modeling workflow according to the group needs, namely aggregating species occurrences and running automated data validation routines, modeling species distributions and processing expert’s feedback on models. Additionally, the core team approves the creation of new groups, enables the publication of models generated by third parties and updates occurrence databases following recommendations provided by the groups.

## Model development workflow

Species distribution hypothesis available in the BioModelos web app are generated either by collaborative development of species distribution models and expert maps or by third parties that independently upload models to the web app (Fig 1).

**Fig 1.**
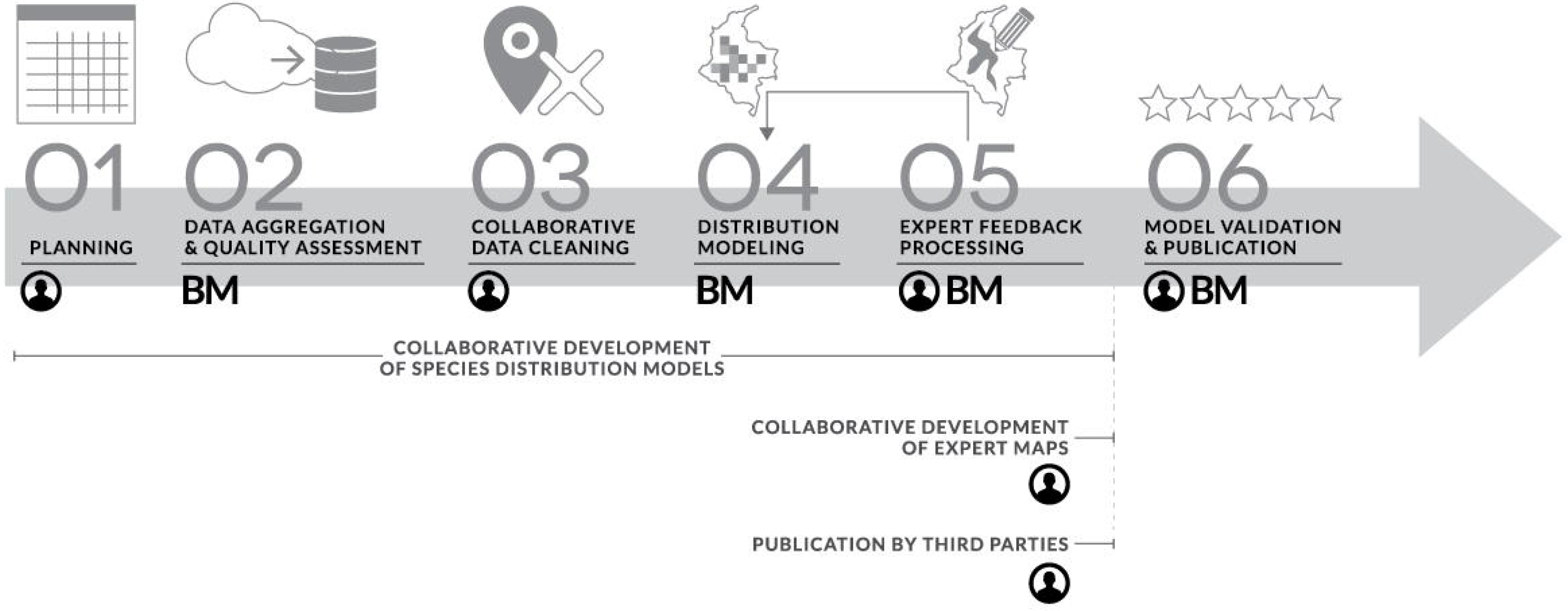
Species modeling workflow. Activities developed by the core team are indicated by BM whereas activities developed by experts are indicated by encircled person logo.

### Collaborative development of species distribution models Planning

In BioModelos, every species is associated to a single group of experts whom are tasked with generating, improving or validating a distribution hypothesis for each species in the group. Experts within a given group inform the core team which species they plan to model collaboratively within a given time frame and provide information that may be relevant for model development, such as previously cleaned occurrence data or not publicly accessible, suggestions of meaningful environmental variables to consider in modeling, among others.

### Data aggregation and quality assessment

Unless the group provides previously cleaned data, the core team aggregates occurrences from multiple data providers (GBIF, eBird, VertNet etc.), either manually or through web services when available. After aggregation and standardization, a series of automated data quality checks are performed (further info in Table 1). Importantly, a permanent unique identifier for each occurrence is generated and original identifiers (e.g. occurrence id, institution, collection code, catalog numbers etc.) are maintained throughout the process so that their provenance may be traced and feedback on data quality may be sent back to data providers whenever mechanisms to that end exist.

**Table 1.**
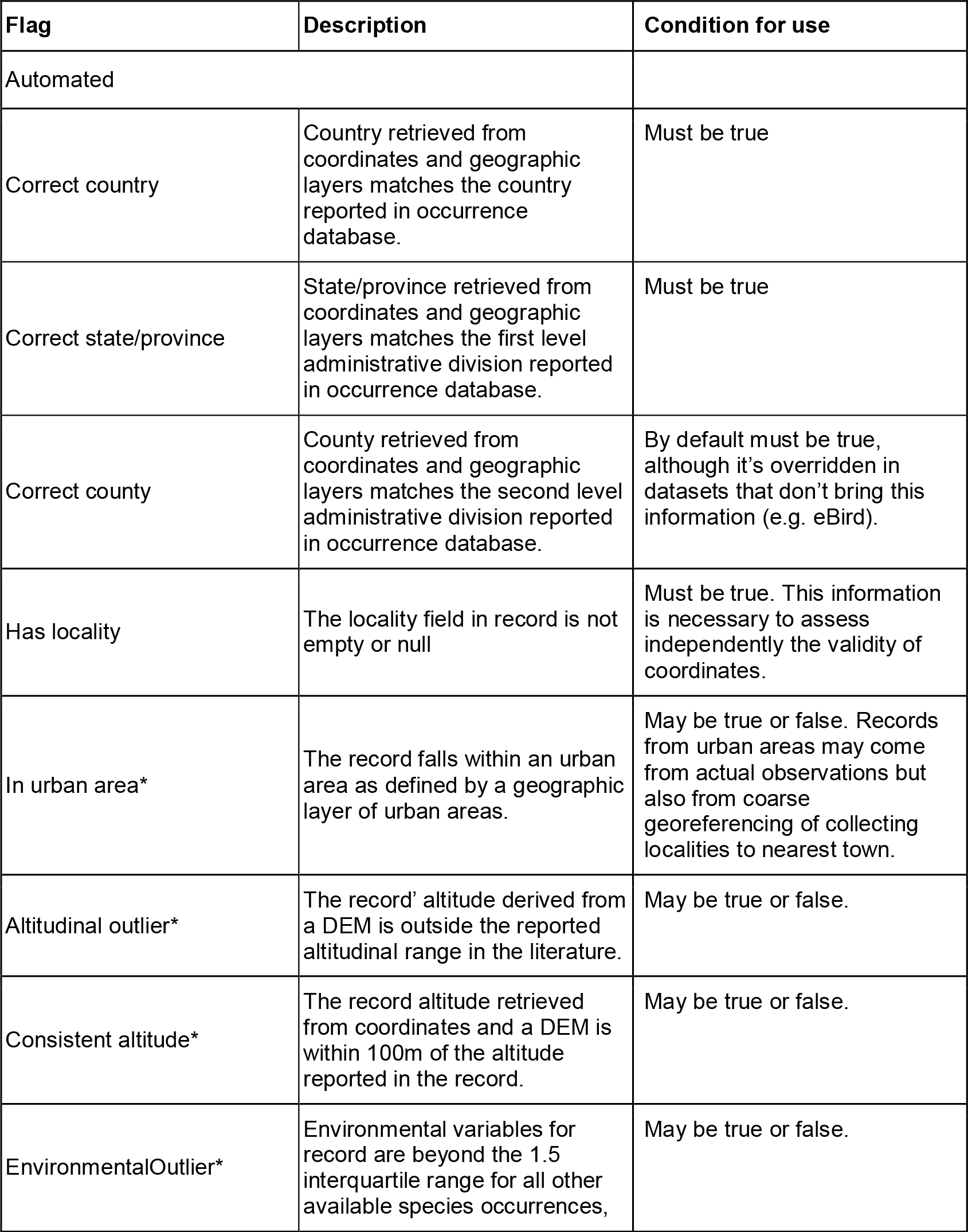
Automated and expert data validation and cleaning procedures in BioModelos. *Records that meet this condition are highlighted in the BioModelos viewer for priority review.

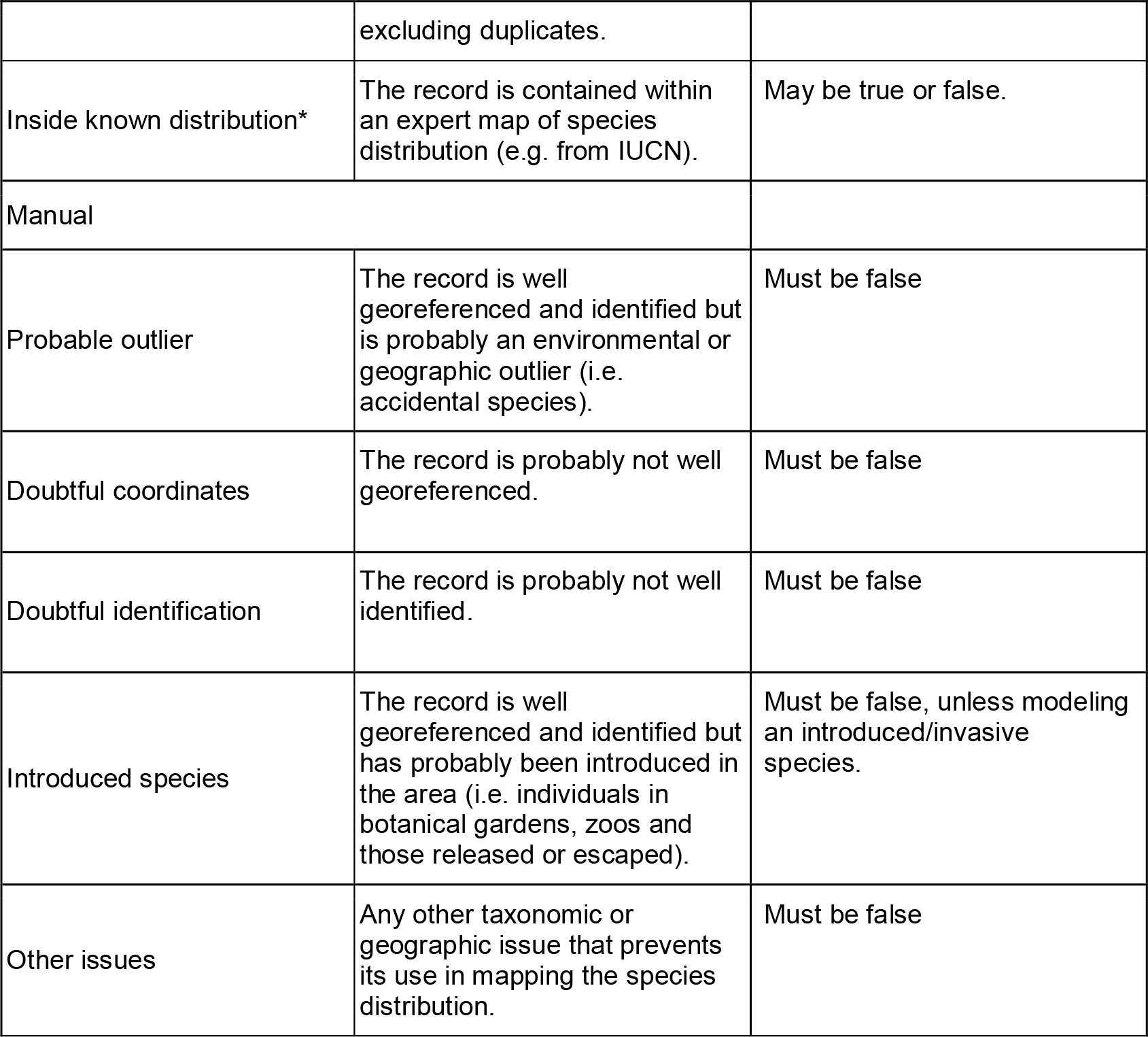

### Collaborative Data Cleaning

Records that pass selected filters based on the automated quality checks become visible on the BioModelos site. The filters implemented for each dataset are variable, depending on the characteristics of each dataset and the amount of records available for a particular group. Published records are further reviewed in BioModelos by experts to identify and flag likely identification and georeferencing errors not detectable in automated checks. Whenever corrections are possible, experts are encouraged to edit identifications or coordinates, while keeping a log of changes in the records database. If a correction is not possible, experts flag records according to the manual categories presented in Table 1.

To inform the development of distribution models, experts are also asked at this step to delineate rough polygons of species’ accessible areas [M sensu 37] using a polygon tool and identify land cover types where species are expected to maintain viable populations by filling out a habitat preferences form.

### Distribution modeling

After occurrence data has been assessed for quality, the BioModelos core team runs species distribution models using occurrences without quality issues and publishes them in BioModelos with status “under development”. As no single modeling workflow is guaranteed to be adequate for all species [26,29], there is considerable variability from group to group in terms of modeling choices, namely, occurrence thinning, variable selection, modeling method, complexity optimization, background extent and background sampling strategy. Therefore, we emphasize that the BioModelos web app acts as a layer through which information from experts is gathered, but once collected a variety of modeling workflows may be implemented. This allows us to update our modeling workflow continuously based on advances in the field independently from the BioModelos web app.

### Expert feedback processing

Models published in the previous step are reviewed by experts whom select through a slider an omission threshold (varying from 0% to 30%) to convert continuous models into binary models. When areas of model overprediction are identified, they may also use a polygon tool to delineate them. Once these inputs are processed by the core team the resulting models are published in the “available hypothesis” box of BioModelos (Fig. 2).

**Fig 2.**
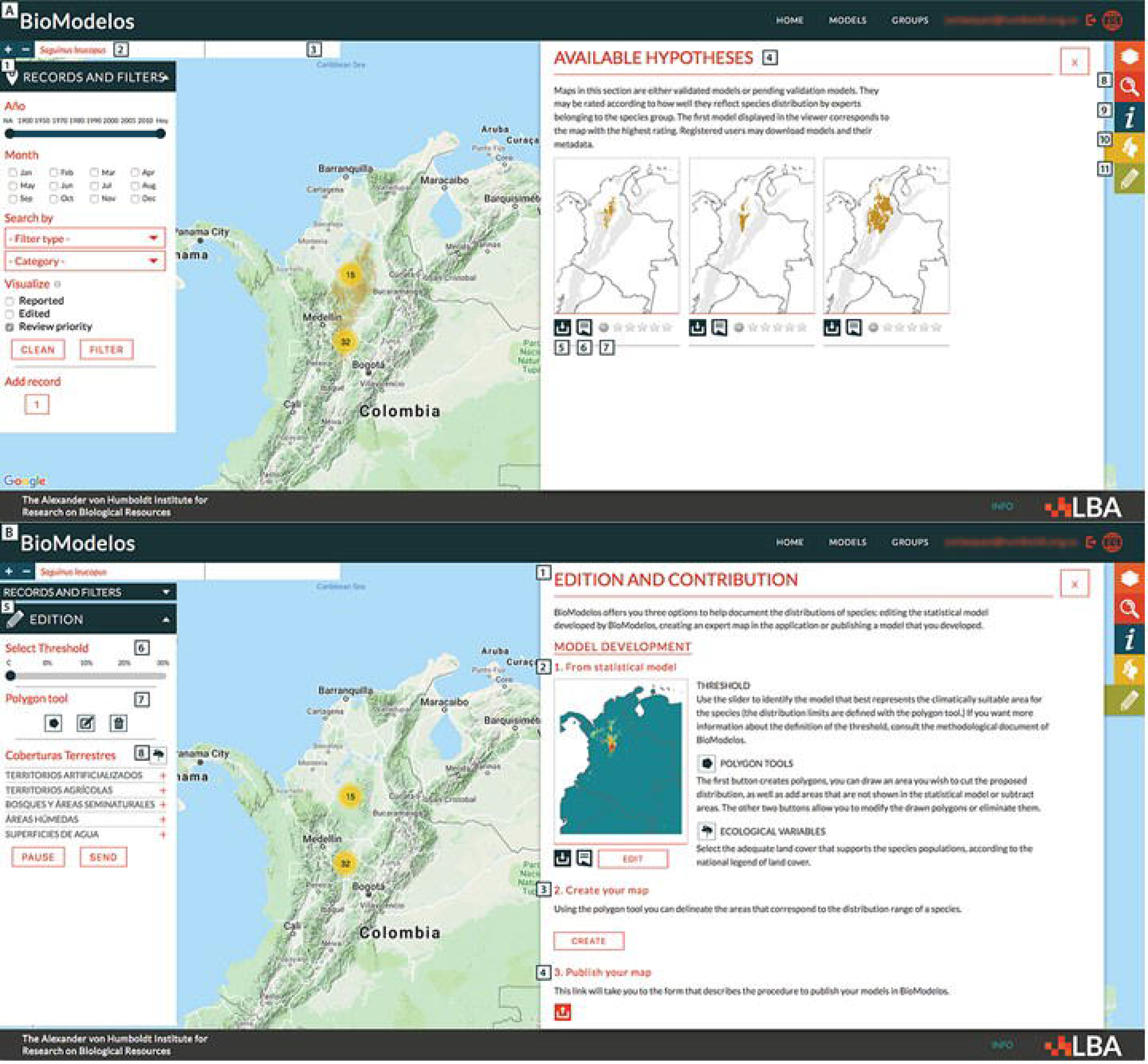
**A. BioModelos geographic viewer subcomponents.** (1) Occurrence filter panel; (2) Species name; (3) Model status; (4) Available hypothesis box; (5)Download link; (6) Metadata link; (7) Score model; (8) Advanced search box; (9) Species fact-sheet box; (10) Available hypothesis box; (11) Generate map and edit model panel. **B. BioModelos model generation and input tools (only viewable for experts working on a species in their task dashboard).** (1) Model generation box; (2) Model input option; (3) Create map option; (4) Publish map option; (5) Model input panel; (6) Threshold slider; (7) polygon tools to delimit accessible areas and identification of areas of model over and underprediction; (8) species habitat preferences formulary.

### Collaborative development of expert maps

For many species without sufficient occurrence data it is not possible to develop a distribution model. In those cases, experts may still use BioModelos to inform the general range limits of a species and their habitat preferences using the “create habitat map” feature (Fig 2.). Further, by choosing this option, experts can provide instructions to further refine them based on geographic features such as barriers (rivers, canyons, watersheds) and elevation. These instructions are processed by the core team to generate species distribution maps that are displayed in the BioModelos viewer in the “available hypothesis” box.

### Publication by third parties

As a rapidly expanding field, a large number of species distribution models are being generated by numerous researchers. Many of these are produced by experts on the species being modeled, and thus represent a valuable resource to other scientists and users. The publication of these models is encouraged in BioModelos, as a way to maintain and facilitate their access. To publish a model in BioModelos the following requirements must be met: (1) occurrence data must have had a data cleaning process to identify and correct georeferencing and identification errors; (2) modeling should follow a minimum of accepted best practices in the field [e.g. optimization of regularization and feature selection in Maxent; 27]; (3) predictions should be restricted to each species’ accessible area. Submissions are for the most part verified by the core team, though in some instances they may be verified by an external reviewer.

## Model validation and publication

An essential feature of BioModelos is the experts’ approval of a distribution hypothesis for a species. This approval is inherently subjective and is made on the basis of experts’ subjective judgement on the biological realism of a model. Since at any point there may be several hypotheses for the distribution of a species (for example a species may have a two published models and one collaborative model), we ask experts in each group to score the models for species in their group, on a qualitative scale from 1 to 5 (1: no credibility, 5: complete credibility). Models with an average score of 3 or higher are approved and flagged as “*validated*”. If no model is approved, experts may either decide to go back and modify their inputs or suggest the development of a new model altogether. The process of expert model evaluation is repeated whenever new distribution hypothesis are generated, taxonomy changes or significantly new occurrence data becomes available. Once a species has a validated model within BioModelos, the core team calculates a number of statistics based of its distribution to aid the assessment of its conservation status and trends which are displayed in the species fact-sheet box (Fig. 2).

All distribution models visible in BioModelos are available for download in GeoTIFF formal at their original resolution and their use and distribution is regulated under Creative Commons 3.0 licenses. Everyone involved in the data cleaning, generation of model inputs and validation of models are recognized as model’ authors. Besides including all author information, our metadata standard for models and occurrences (Appendix 1), contains all relevant information pertaining to data sources, model development (including links to modeling logs in GitHub) and performance statistics.

## Web application architecture and components

BioModelos has been developed as an open source web application composed of four main components over three layers: a website database, a contents database, a RESTful API and the web application. Each component runs independently on its own container deployed using Docker (https://www.docker.com/), giving an increased granularity in the control and maintenance of the components. At its core, BioModelos relies on a content and a website database. The content database was developed following a non-relational scheme in MongoDB (https://www.mongodb.com) and it includes the collections “species” (keeps the taxonomic backbone and species’ ancillary information), “records” (occurrence data and quality assessment information) and “models” (model metadata and distribution derived statistics). The website database was developed using a relational scheme via PostgreSQL (https://www.postgresql.org/) and it stores all relevant information about the interaction of the users within the website, such as users, groups, user created layers, ratings, tasks, publications, and downloads among others. The website database is directly connected to the BioModelos front-end while the content database is accessible through a RESTful API developed in Javascript. This architecture allows BioModelos to scale, grow and distribute each database independently as well as to store user interactions on the website privately while opening the BioModelos contents to third-party applications.

The front-end of BioModelos was developed using the Ruby on Rails (https://rubyonrails.org/) framework. Its main parts are a search engine, a social network component and a geographic viewer. The search engine allows users to find species distributions by entering their scientific name or sets of species based on attributes using the advanced search functions. The social network component comprises expert’s public profiles and group profiles and allows experts interaction using built in messaging tools, monitoring of progress in completing particular modeling tasks using the task dashboard and approval for the admission of new group members. Lastly, the geographic viewer contains the following subcomponents (Fig. 2; further details in Appendix 1):

> Data cleaning panel: contains tools to add occurrences to the “records” collection, flag occurrences with identification or georeferencing issues and edit certain fields for correction or standardization purposes.
>
> Model generation box: contains tools that may be used to either provide input for the development of a model, create an expert map or to publish a model developed independently by a researcher. Whenever the first option is chosen, a set of model input tools becomes available on a side panel which allows experts to provide inputs for model pre and post-processing. Pre-processing, experts may delineate polygons of accessible areas using a polygon tool and identify land cover types where species are expected to maintain viable populations. Post-processing, experts may use a slider to select an omission threshold and use a polygon tool to identify areas of model over and underprediction.
>
> Available hypothesis box: all available hypothesis for the distribution of a species, use licenses and links to their metadata are displayed in this section. Also, here is where experts may rate distribution hypothesis. Download links are provided in this section to registered users.
>
> Species fact-sheet box: contains statistics based on validated species models (e.g. extent of occurrence, representation in protected areas, forest loss within range) and links to the occurrence metadata on which the model is based.

## Implementation of BioModelos in Colombia

The BioModelos software and network have both been under simultaneous development since 2013. During this time we have conducted 15 workshops with experts that have been essential to consolidate expert groups as well as to gather feedback to enhance user experience in the web app and improve the modeling workflow. Currently, out of 1052 registered users, there are 475 experts associated to 20 expert groups (Table 2), which in turn are managed by 34 moderators. Collectively, these experts are tasked with contributing in the development of 980 SDMs. Additionally, 17 expert maps and 216 models have been published since the model publishing feature was implemented in January 2017.

**Table 2.** Summary of the expert network of BioModelos and status of models by groups. Developing: statistical model generated after the process of data aggregation a cleaning;

| Group name | Number of experts | Species in group | Model development status (spp.) |  |  |
| --- | --- | --- | --- | --- | --- |
|  |  |  | Developing | Pending validation | Valid |
| Bees of Colombia | 12 | 16 |  |  |  |
| Aquatic birds of Colombia | 48 | 80 | 2 |  | 14 |
| Birds of Colombia | 36 | 277 | 49 | 156 | 4 |
| Beetles | 18 | 4 | 1 |  |  |
| Introduced fauna | 24 | 21 | 17 |  | 1 |
| Plants of paramo | 21 | 212 | 212 |  |  |
| Herps of Colombia | 60 | 67 | 60 | 1 | 2 |
| Introduced plants of Colombia | 33 | 32 | 31 |  |  |
| Dragonflies Colombia | 5 | 13 | 11 |  |  |
| Magnolias of Colombia | 17 | 34 | 17 | 13 |  |
| Mammals of Colombia | 81 | 52 | 6 | 1 |  |
| Orchids of Colombia | 25 | 5 |  |  |  |
| Palms | 8 | 51 | 46 |  |  |
| Freshwater fishes | 10 | 1 | 1 |  |  |
| Carnivorous plants of Colombia | 4 | 4 | 1 |  |  |
| Plants of dry forest | 28 | 53 | 48 |  |  |
| Primates of Colombia | 28 | 38 |  | 1 | 37 |
| Zamias of Colombia | 17 | 20 |  |  | 20 |
| **TOTAL** | **475** | **980** | **502** | **172** | **78** |

We began requesting user registration to download models since August 2017. Ever since, 430 downloads have been made in BioModelos, 33% of them academic research, 32% for educational activities and the remaining for applied research, environmental consulting, bioprospecting and other activities. Validated models have allowed Humboldt Institute (BioModelos’ host institution in Colombia) to support conservation decision making by informing biodiversity loss compensation plans [38], extinction risk assessment of species [39] and support the generation of official cartography for Colombia, like the Colombian ecosystems map [40].

## Discussion

We presented BioModelos, an approach to facilitate collaboration between experts (e.g. field biologists, ecologists, taxonomists, biogeographers and modelers) to generate publicly available information on species distribution mediated by a core team and a web app. By involving experts in the development of models, we aim to fill the gaps in primary biodiversity data and assess the biological realism of model predictions by eliciting experts’ opinion on species distribution as well as to avoid the prevalent duplication of efforts in data cleaning and modeling (Anderson et al. 2016). Both of these features are necessary to advance faster towards the amelioration of the Wallacean shortfall as well as to further the use of SDMs to generate EBV that inform conservation decision making processes [15].

Species distribution modeling is still a very dynamic field in which novel methods and recommendations arise frequently. By keeping the expert-opinion data gathering process independent from the modeling process, we have been able to implement multiple modeling workflows with little impact on the design of the BioModelos GUI or user’s experience, keeping the app maintenance costs low. This feature will allow us to continue to refine our modeling workflow for example by using methods that formally incorporate expert opinion in model development [41] and integrating established semi-automated modeling workflows such as Wallace [42].

The difficulty of evaluating the accuracy of species distribution models in presence-only models has long been recognized and discussed [43,44]. Simply put, traditional performance metrics (e.g. AUC, TSS) of models built based on presence-only data may only tell us how well a model prediction discriminates presences from arbitrary pseudo-absences and their value and statistical significance depends on how those pseudo-absences are drawn. Therefore, these metrics are a measure of relative performance not suitable for comparison among species and of difficult interpretation for interested users of model predictions without a modeling background. Another important feature of BioModelos is that besides providing standard measures of model performance for each published model it also encourages the subjective evaluation of models by experts. This qualitative evaluation together with authorship information on the experts that validate the model may help end-users to decide whether to use a model or not in a particular application.

An important challenge of the implementation of BioModelos in Colombia has been to motivate the autonomous completion of modeling agendas by experts’ groups. Thus far, the most successful collaborative modeling experiences have been with groups that need BioModelos outputs (i.e. models and species fact-sheets) for species risk assessments [waterbirds; 39] or action plans (Cycads, Magnoliaceae and Primates; in progress). In the near future we aim to bring together more experts focusing on species of conservation concern as those will likely benefit more from having high quality distribution information. Also, the publication of models developed by third-parties has contributed an important proportion of the models available in BioModelos and we continue to encourage model publication by contacting modelers identified through publications and scientific events. However, for the remaining species we are still in the process of devising incentives to increase the participation of experts in BioModelos, such as the development of electronic publications that are formally recognized as research products in academic performance reviews.

The BioModelos web app is open source (https://github.com/LBAB-Humboldt/BioModelos.v2) and free to use by any interested party. However, due to the technical requirements for its installation and maintenance, the collaborative nature of the BioModelos approach and the need of a core team consisting at least of a modeler and a network manager for its implementation, it is best suited for national level implementations hosted by research institutions. Although this geographical scale may seem arbitrary as species are not limited *per se* by national boundaries, it makes sense considering that many uses of models for conservation decision making take place at national and subnational levels and that experts usually confine their expertise to countries due to accessibility, funding restrictions, easiness in obtaining research permits etc. Hence, by implementing BioModelos at a national scale, we contribute both to increase occurrence data fitness for use in distribution modeling, potentially aiding global modeling initiatives once mechanisms to collect error reports through web services are implemented in data providers such as GBIF [45] and to the consolidation of a global coordinated monitoring system of species distributions through National Biodiversity Observation Networks [46], to make EBV’s genuinely global [47].

## Acknowledgements

We are grateful for the feedback provided in the past five years by the numerous researchers that have attended our workshops and the institutions they represent: Asociación Colombiana de Ornitología, Calidris, Asociación Bogotana de Ornitología, Asociación Colombiana de Herpetología, ASICTIOS, Sociedad Colombiana de Mastozoología, Universidad Nacional, Universidad Javeriana, Universidad de Antioquia, Universidad del Valle, among others. Their input has been critical to enhance the contents of BioModelos as well as designing a user friendly web app. Particularly, we want to thank Cristina López Gallego, Luis Francisco Sanchez and Nicolás Urbina for their advice on developing a feasible collaborative species modeling workflow and their energy in moving forward the BioModelos agenda in the groups they moderate. This project would not have been possible without the institutional backing of BioModelos provided by Instituto Humboldt, particularly by Brigitte Baptiste, Hernando García, Germán Andrade, Juan Carlos Bello, Jose Ochoa, Johanna Galvis and Cristina Ruiz. Carolina Bello, Valentina Grajales, Lina Estupiñán and Ricardo Bastidas provided technical input for the development of BioModelos at several stages. This manuscript was greatly enhanced by comments provided by Robert Anderson.

## Author Contributions

Conceived the project: JVT MCLM. Designed methodology: JVT MCLM MHOR IG. Software development: DLL. Visualization: CG. Wrote the paper: JVT, MCLM, MHO, DL.

## Supporting Information

**Table S1**. BioModelos’ main features

